# Identifying new cancer genes based on the integration of annotated gene sets via hypergraph neural networks

**DOI:** 10.1101/2024.01.22.576645

**Authors:** Chao Deng, Hong-Dong Li, Li-Shen Zhang, Yi-Wei Liu, Yaohang Li, Jianxin Wang

## Abstract

**Motivation:** Identifying cancer genes remains a significant challenge in cancer genomics research. Annotated gene sets encode functional associations among multiple genes, and cancer genes have been shown to cluster in hallmark signaling pathways and biological processes. The knowledge of annotated gene sets is critical for discovering cancer genes but remains to be fully exploited.

**Results:** Here, we present the DIsease-Specific Hypergraph neural network (DISHyper), a hypergraph-based computational method that integrates the knowledge from multiple types of annotated gene sets to predict cancer genes. First, our benchmark results demonstrate that DISHyper outperforms the existing state-of-the-art methods and highlight the advantages of employing hypergraphs for representing annotated gene sets. Second, we validate the accuracy of DISHyper-predicted cancer genes using functional validation results and multiple independent functional genomics data. Third, our model predicts 44 novel cancer genes, and subsequent analysis shows their significant associations with multiple types of cancers. Overall, our study provides a new perspective for discovering cancer genes and reveals previously undiscovered cancer genes.

**Availability:** DISHyper is freely available for download at https://github.com/genemine/DISHyper.

## 1. Introduction

Cancer is a set of complex and highly genetically heterogeneous diseases (Lawrence et al., 2013). A key goal in cancer genomic research is to identify cancer genes that play a causal driving role in the development and progression of tumors (Garraway and Lander, 2013; Alexandrov et al., 2013; Vogelstein et al., 2013). The identification of cancer genes is crucial for the study of tumor pathogenesis, early screening, and the development of precision oncology (Garraway and Lander, 2013; Alexandrov et al., 2013; Vogelstein et al., 2013). Although many initiatives such as the Network of Cancer Genes (NCG) (Repana et al., 2019) and the COSMIC Cancer Gene Census (CGC) (Sondka et al., 2018) have been used to annotate some cancer genes, the current catalog of known cancer genes is incomplete. Accurate identification of cancer genes from many candidate genes remains a critical challenge.

In the last decade, large-scale cancer sequencing projects such as The Cancer Genome Atlas (TCGA) (Weinstein et al., 2013) and Pan-Cancer Analysis of Whole Genomes (PCAWG) (Pan-Cancer Analysis of Whole Genomes Consortium, 2020) have published genomic and transcriptomic data from tens of thousands of tumor samples. These large-scale cancer sample data have largely contributed to the development of computational methods for cancer gene identification. Early approaches for cancer gene prediction focused on finding genes that have significantly different mutation rates from the background frequency distribution, such as MutSigCV (Lawrence et al., 2013). Meanwhile, the 20/20+ method proposed finding genes with similar mutation patterns to known cancer genes by integrating multiple gene mutation feature through machine-learning models (Tokheim et al., 2016). Moreover, some approaches are proposed to identify cancer genes by integrating features from the genome, transcriptome, and proteome. For example, DORGE proposed integrating epigenetic and mutational features based on machine learning models to identify oncogenes and tumor suppressor genes (Lyu et al., 2020). Biological network-based cancer gene prediction methods have been extensively studied in recent years. HotNet2 (Reyna et al., 2018) used heat diffusion models to detect cancer gene modules with mutational features. EMOGI (Schulte-Sasse et al., 2021) integrates multi-omics data for multiple cancers and protein-protein interaction (PPI) networks via graph convolutional networks (GCN) (Kipf and Welling, 2017) to learn local feature patterns of cancer genes.

Cancer genes have been shown to cluster in a small number of biological processes, hallmark signaling pathways, and interacting subnetworks (Creixell et al., 2015; Reyna et al., 2020). Currently, research on cancer gene prediction methods focuses on the binary associations of cancer genes in biological networks, while ignoring the functional associations of cancer genes in annotated gene sets such as biological processes and signaling pathways. The annotated gene set encodes a group of genes with functional associations (Liberzon et al., 2015). The functional associations in annotation gene sets are higher-order associations among multiple genes, and the higher-order functional association information is more intuitive and accurate in explaining the disease mechanism than single genes (Luo and Mao, 2021; Luo, 2022). In addition, cancer driver mutations can lead to oncogenic properties of cells by altering the activities of hallmark biological processes and pathways (Creixell et al., 2015; Reyna et al., 2020). Therefore, compared with the binary association in biological networks, the higher-order association information in annotated gene sets provides a more comprehensive characterization of the functional association patterns of cancer genes and help us identify cancer genes more accurately. To achieve this goal, it is necessary to develop a computational framework that can integrate knowledge from different types of annotated gene sets and exploit their higher-order association information to identify cancer genes. Currently, a few computational approaches have used annotated gene set data, these approaches typically represent annotated gene sets as graph structures or encode them directly as vectors (Luo et al., 2019; Valdeolivas et al., 2019; Althubaiti et al., 2019). Although graph or network structures have been widely used in computational biology to represent binary associations between biological entities, they are unable to represent higher-order gene associations in annotated gene sets. Therefore, although the knowledge of annotated gene sets is crucial for cancer gene prediction, using the knowledge to identify cancer genes faces the following problems: (i) how to represent the higher-order association among multiple genes in annotated gene sets, (ii) how to integrate different types of annotated gene sets such as gene ontology (GO), signaling pathways, and human phenotype ontology (HPO), (iii) how to make full use of the higher-order functional association information to identify cancer genes.

To tackle these problems, we introduce DISHyper, a novel method to identify cancer genes based on the annotated gene sets and hypergraph neural networks (HGNN). In DISHyper, we first represent and integrate different types of annotated gene sets using the hypergraph structure. Each gene is a node in the hypergraph, and each annotated gene set is a hyperedge. Hypergraph provides a natural way to represent the higher-order association relationships between multiple genes in annotated gene sets. Then, to fully utilize the higher-order association information among multiple genes in the annotated gene sets, we use HGNN to train the model.

The HGNN extracts higher-order gene association information and local topological information from the annotated gene set and generates the feature representation of the gene through the two-phase message-passing process. We use this feature representation to prioritize cancer genes. In addition, we propose the disease-specific hyperedge weighting module and the hypergraph residual learning module based on HGNN. We use these two modules to weight the knowledge for each annotated gene set and enhance the expressive power of the model.

We evaluate the performance of DISHyper and predict new cancer genes based on pan-cancer data. In our benchmark experiments, DISHyper outperforms the other state-of-the-art methods and has a significant performance improvement over the other methods. Through comprehensive ablation studies, we also show the effectiveness of DISHyper in integrating multiple types of annotated gene sets and the importance of using disease-specific weighting modules and hypergraph residual learning modules. We conduct a comprehensive assessment of DISHyper-predicted cancer genes using multiple analysis methods and data. Firstly, we extensively analyze and evaluate the DISHyper-predicted top-rank 200 cancer genes (PCG) using functional validation experiment results and independent functional genomic data. Then, we perform a comprehensive enrichment analysis of DISHyper-predicted cancer genes. Finally, we perform further evaluation for our predicted 44 novel cancer genes (novelCG) based on RNA-Seq, DNA methylation, and clinical data from tumor samples in the TCGA study. Our analysis shows that the 44 novel cancer genes are closely associated with one or more cancers. Moreover, We illustrate the DISHyper prediction process through a case study of *WNT5A*. To the best of our knowledge, DISHyper is the first hypergraph-based cancer gene prediction computational method that utilizes higher-order functional association information among multiple genes to identify cancer genes.

## 2. Materials and methods

### 2.1 Datasets and processing

We collect pan-cancer data to evaluate model performance and predict new cancer genes. We use the same list of known cancer genes and non-cancer genes as in EMOGI (Schulte-Sasse et al., 2021) to evaluate model performance and predict new cancer genes. Specifically, we collect known cancer driver genes from the NCG (Repana et al., 2019) (v6.0), COSMIC CGC (Sondka et al., 2018) (v91), and DigSEE databases (Kim et al., 2013) as positive samples. The negative samples are the remaining genes after recursively excluding the NCG, COSMIC CGC, OMIM database, DigSEE database, and KEGG cancer pathway gene set. Thus, our pan-cancer gene data consists of 796 positive samples and 2,187 negative samples.

We collect multiple types of annotated gene sets from the Molecular Signatures Database (MSigDB) (Liberzon et al., 2015). The MSigDB (v7.4) database contains multiple types of annotated gene set data and classifies them into nine categories. We use C2 and C5 gene sets from the MSigDB database, which contains signaling pathway data from expert databases such as BioCarta (Rouillard et al., 2016), Reactome (Fabregat et al., 2018), and KEGG (Kanehisa and Goto, 2000), and ontology gene sets from GO (Gene Ontology Consortium, 2004) and HPO (Köhler et al., 2021). To accurately assess the model performance, we eliminate the annotated gene sets directly annotated with cancer during model training and testing. We collect a total of 20,647 annotated gene sets, encompassing functional associations among 17,442 genes.

### 2.2. Framework of DISHyper

DISHyper is a novel hypergraph-based cancer gene prediction method. The workflow of DISHyper is shown in Figure 1 and can be summarized into two components: (1) Construction of disease-specific hypergraph. We represent and integrate different types of annotated gene sets with hypergraph structure, and perform disease-specific hyperedge weighting using priori information on cancer to construct disease-specific hypergraph. (2) Higher-order association information learning via hypergraph residual neural networks. We use the construct disease-specific hypergraph and the initial feature matrix **X** of genes as the input to train the cancer gene prediction model. We first perform a nonlinear transformation on the feature matrix, and then use three hypergraph residual learning modules to extract higher-order association information of genes and generate new feature representations for genes. Finally, we predict the degree of association between each gene and cancer.

**Fig. 1:**
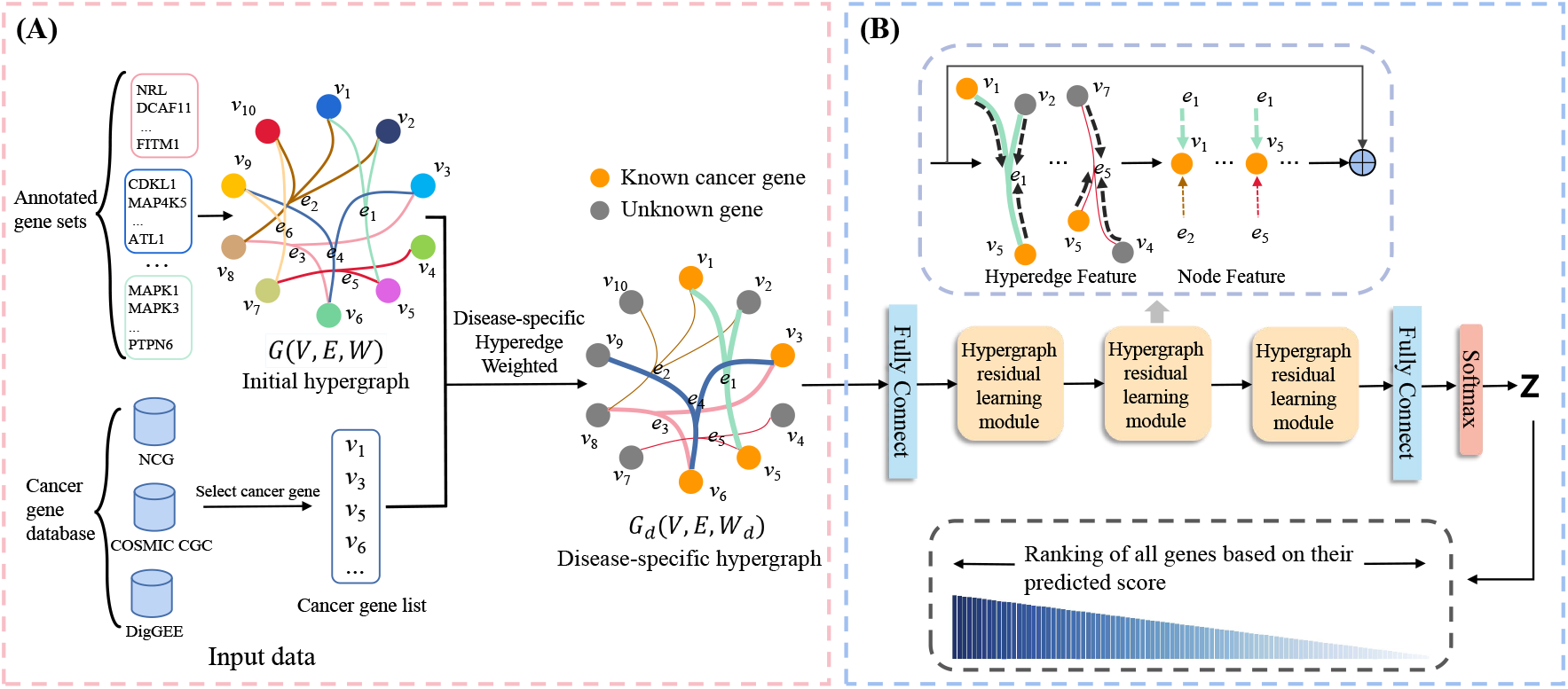
Illustration of DISHyper. **(A)** Construction of the disease-specific hypergraph. We collect the annotated gene set data from MSigDB and construct the initial hypergraph *G*(*V, E*, **W**). We then perform disease-specific hyperedge weighting with known cancer genes as prior knowledge and construct the disease-specific hypergraph *G*_*d*_(*V, E*, **W**_**d**_). **(B)** Higher-order association information learning via hypergraph residual neural networks. We take the disease-specific hypergraph *G*_*d*_(*V, E*, **W**_**d**_) and the initial feature matrix **X** as input of the model. We first perform a nonlinear transformation on the feature matrix **X** by one fully connected layer and activation function. Then, we use the hypergraph residual learning module to extract local topology and higher-order association information of genes in the disease-specific hypergraph *G*_*d*_ and predict the risk score **Z** for each gene.

#### 2.2.1 Construction of the disease-specific hypergraph

We use DISHyper to integrate knowledge from different types of annotated gene sets. Each annotated gene set represents a set of genes with functional associations. For example, a signaling pathway indicates that multiple genes are participating in a particular signaling process. To accurately describe the complex associations of genes in annotated gene sets and fully utilize the higher-order association information in annotated gene sets, DISHyper uses the hypergraph structure to represent and integrate different types of annotated gene sets. We define the initial hypergraph constructed using annotated gene sets as *G*(*V, E*, **W**), the node *V* = {*v*_1_, *v*_2_, *v*_3_, …, *v*_*n*_} in the hypergraph represents *n* genes, and the hyperedge *E* = {*e*_1_, *e*_2_, *e*_3_, …, *e*_*m*_} represents *m* annotated gene sets. The **W** denote the diagonal matrix of the hyperedge weights, i.e., *diag*(**W**) = [*w*(*e*_1_), *w*(*e*_2_), …, *w*(*e*_*m*_)]. **W** is generally an identity matrix. For the incidence matrix **H** ∈ ℜ^*n*×*m*^ of the hypergraph, we define that if the *a*-th gene *v*_*a*_ belongs to the *b*-th annotated gene set *e*_*b*_, then **H**(*v*_*a*_, *e*_*b*_) = 1, otherwise 0.

In the node classification task of the hypergraph, the weight assigned to a hyperedge reflects its significance in the classification process. The weight matrix **W** in the initial hypergraph *G* is the identity matrix. If the initial hypergraph *G* is used to train the model directly, then each annotated gene set (hyperedge) has the same importance for that cancer gene prediction task. However, the different annotated gene sets obviously have different degrees of impact on the cancer. For example, the biological processes associated with cell proliferation and cell metastasis are more closely associated with cancer. Therefore, those annotated gene sets with more significant associations with the process of tumor development should be assigned greater hyperedge weights. Considering the association specificity of cancer to different annotated gene sets, we propose a disease-specific hyperedge weighting module. We use the known cancer genes as the prior information of cancer and calculate the proportion of known cancer genes in each annotated gene set as hyperedge weights. The weight of the *i*-th hyperedge is defined as follows:

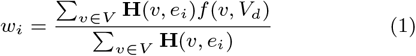

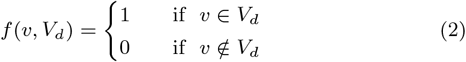

where *w*_*i*_ denotes the weight of the *i*-th (*i* ∈ [1, *m*]) hyperedge, and *V*_*d*_ denotes the set of known cancer genes. The *f* (*v, V*_*d*_) is used to indicate whether gene *v* belongs to *V*_*d*_. We define the disease-specific hypergraph as *G*_*d*_(*V, E*, **W**_**d**_), where **W**_**d**_ is the weighted hyperedge weight matrix. The disease-specific hypergraph *G*_*d*_ contains the disease-specific functional association between cancer and annotated gene sets. For annotated gene sets with a larger proportion of known cancer genes, we assign proportionally greater hyperedge weights to them because these annotated gene sets are more likely to be relevant to the cancer gene prediction task. It should be emphasized that, in benchmarking, we use only the positive genes in the train set for disease-specific hyperedge weighting, and the positive genes in the test set are excluded in this process.

#### 2.2.2 Higher-order association information learning via hypergraph residual neural networks

HGNN is a semi-supervised hypergraph node classification method (Feng et al., 2019). We model the disease-specific hypergraphs *G*_*d*_ through HGNN and extract complex association information of genes in annotated gene sets. Both HGNN and GCN aggregate the neighbor information of nodes through the message-passing mechanism and generate the new feature representation containing neighbor information for each node. Stacking multiple layers of GCN means that each node will aggregate multi-order neighbor information. When the GCNs are deep, the features of the nodes tend to be consistent and difficult to distinguish. HGNN also has the risk of over-smoothing, which limits the expressiveness of HGNN and makes the model difficult to converge. At the same time, we also find in the experiment that if two nodes have similar neighbors in the hypergraph, the feature representations of the nodes must be the same after HGNN, even if their initial features and labels are very different. This phenomenon will lead to a loss of specificity in the feature representations of the nodes learned by HGNN and limits the expressive power of HGNN.

Aiming at the problems in HGNN, inspired by ResNet (He et al., 2016) and deepGCN (Li et al., 2019), we develop the hypergraph residual learning module. The formula is defined as follows:

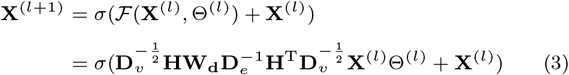

where the **X**^(*l*+1)^ and **X**^(*l*)^ represent the output of the *l* + 1 layer and the *l* layer. We define the degree of the node *v*_*a*_ is *d*(*v*_*a*_) = Σ_*e*∈*E*_ *w*(*e*)**H**(*v*_*a*_, *e*) and the degree of the hyperedge *e*_*b*_ is *δ*(*e*_*b*_) = Σ_*v*∈*V*_ **H**(*v, e*_*b*_). The **D**_**v**_ ∈ ℜ^*n*×*n*^ and **D**_**e**_ ∈ ℜ^*m*×*m*^ denote the diagonal matrices of the node degrees and the hyperedge degrees, respectively. Θ^(*l*)^ is a trainable weight matrix on the *l*-th layer and *σ* denotes the non-linear activation function. The hypergraph residual learning module can be regarded as adding the input **X**^(*l*)^ to the output of the hypergraph convolutional layer. In theory, if the multi-order neighbor information is not needed, the hypergraph residual learning module can make Θ approach **0**, thereby converting the hypergraph convolutional layer into an identity map (**X**^(*l*+1)^ = **X**^(*l*)^), which can reduce the risk of over-smoothing, and accelerate model convergence. We use multi-layer hypergraph residual learning modules to extract the local topology information in the disease-specific hypergraph and generate the new feature representation for each gene.

Finally, DISHyper inputs the gene feature matrix to the fully connected layer and uses the softmax layer to get the prediction result of the model, the formula is as follows:

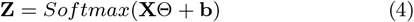

where **Z** ∈ ℜ^*N* ×2^ represents the probability output by DISHyper for each gene being a neutral gene and a cancer gene, and we utilize the latter as the predicted risk score of each gene.

## 3. Results

### 3.1. DISHyper outperforms existing cancer gene prediction methods

To demonstrate the superiority of DISHyper, we conduct a comparison with four state-of-the-art cancer gene prediction methods, namely 20/20+, DORGE, EMOGI, and MTGCN. Furthermore, to emphasize the advantages and necessity of employing the hypergraph for representing annotated gene sets, we compare DISHyper with two graph-based representation methods, including GCN and GCNII. (1) 20/20+ (Tokheim et al., 2016) designed a series of mutation-based feature-trained random forest models to find genes with the same mutation pattern as known cancer genes. (2) DORGE (Lyu et al., 2020) combined mutation data with epigenetic data and added epigenetic features such as methylation, histone modifications, and super-enhancer percentages on the basis of 20/20+. (3) EMOGI (Schulte-Sasse et al., 2021) proposed integrating multi-omics features based on the graph convolution network to identify new cancer genes by learning local neighborhood features of cancer genes. (4) MTGCN (Peng et al., 2022) added the linkage prediction auxiliary task to the PPI network on the basis of EMOGI and optimized the learning of gene features through this multi-task learning framework. (5) GCN: We utilize graph structure to represent annotated gene sets and apply GCN for gene feature extraction. Specifically, we consider genes as nodes, represent each annotated gene set as a complete subgraph, and combine all complete subgraphs into a graph. We use this graph to represent all annotated gene sets and as input for the GCN. (6) GCNII: This variant incorporates initial residual and identity mapping into the GCN (Chen et al., 2020), which offers enhanced model expressiveness and utilizes the same inputs as GCN. To ensure the fairness of comparison, the same positive and negative samples are used in the comparison experiments. For each comparison method and our approach, we conducted multiple times of stratified five-fold cross-validation to assess model performance. The area under the receiver operating characteristic curve (AUROC) and the area under the precision-recall curve (AUPRC) as evaluation metrics for the model performance.

As shown in Figure 2A, DISHyper achieves better performance in comparison with these methods. By taking advantage of hypergraph learning, DISHyper significantly improves in AUROC and AUPRC compared to the two advanced network-based methods, EMOGI and MTGCN. Both DISHyper and these two network-based approaches are essentially finding genes that have similar functions or association patterns to known cancer genes. The results suggest that compared with the binary association in biological networks, the higher-order association among multiple genes in the annotated gene set can capture the complex association patterns among genes more precisely. Compared with 20/20+ and DORGE, two methods based on mutational and epigenetic features, DISHyper shows a 3.6% and 4.8% improvement in AUROC and AUPRC. These manual feature-based methods rely on individual experience and expertise, but the annotated gene set contains multiple expert and domain knowledge from different sources. The benchmark experiment results show that DISHyper can identify cancer genes more accurately than existing advanced methods. Compared with GCN and GCNII, DISHyper shows a 2.4% and 4.5% improvement in AUROC and AUPRC. This is due to the fact that certain crucial information is lost in graph-based representation methods, which ignore higher-order associations among multiple genes within the gene sets. The result implies that hypergraphs can provide a more comprehensive representation of the complex associations in annotated gene sets.

**Fig. 2:**
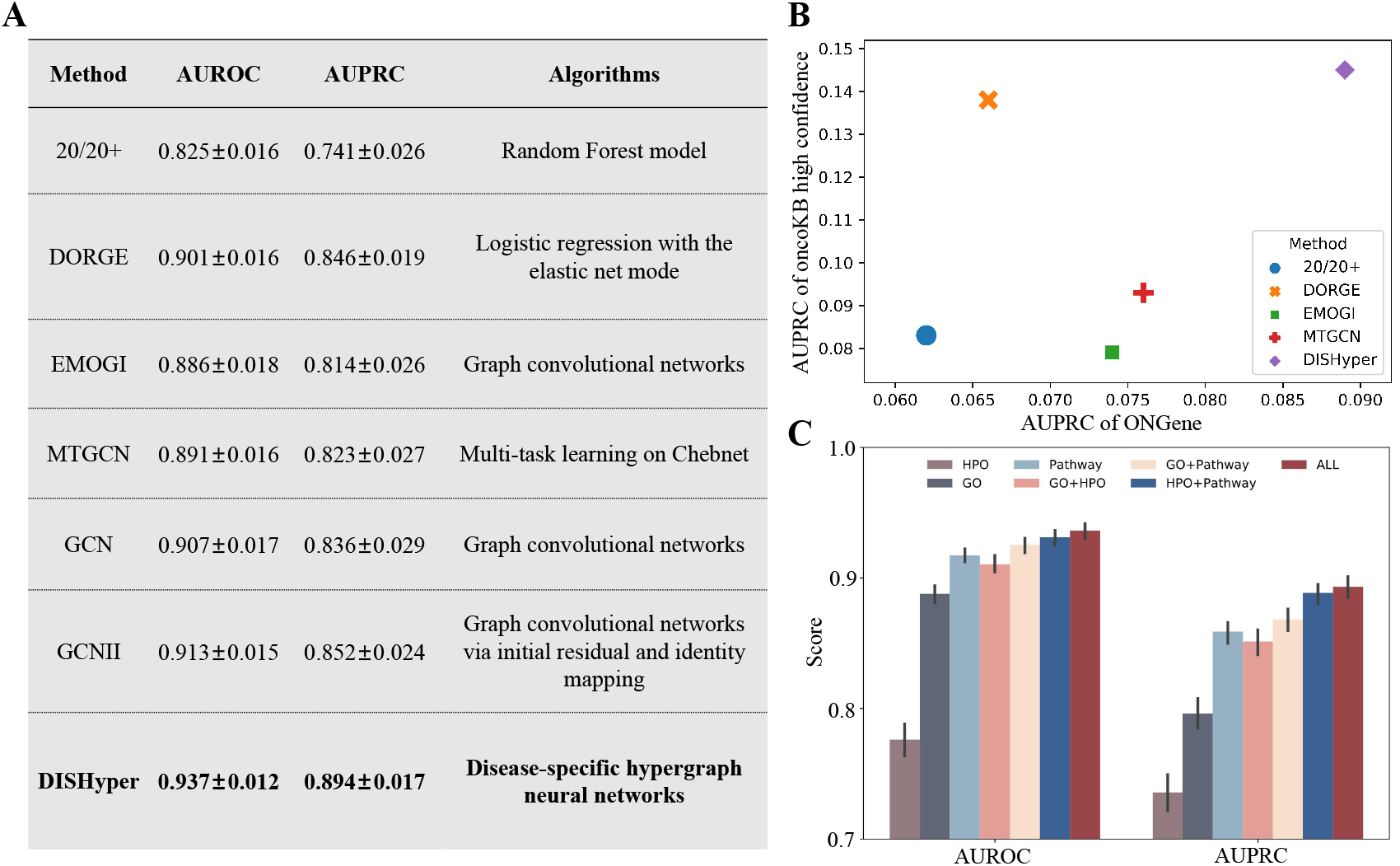
Benchmarking results of DISHyper. **(A)** Comparative experimental results. We compare DISHyper with four state-of-the-art cancer gene prediction methods and two graph-based annotated gene sets representation methods. **(B)** Performance comparison of DISHyper and other methods on two different independent test sets. **(C)** The data ablation analysis results of DISHyper. We analyze the performance of DISHyper using different types of annotated gene sets and their combinations. The error bars in the figure represent 95% confidence intervals.

To further evaluate the generalization performance of DISHyper and its ability to correctly identify cancer genes, we use two sets of curated cancer genes as independent test sets to assess whether the predictions of DISHyper would be biased toward specific cancer genes. The two independent test sets we used are from the OncoKB (Chakravarty et al., 2017) database and the ONGene (Liu et al., 2017) database, with 313 and 382 cancer genes remaining after screening, respectively. To calculate the AUPRC of the model on these two independent test sets, we treat the genes in the independent test set as true positives and all other genes not included in the independent test set as false positives, resulting in much lower AUPRC values for all methods. Figure 2B shows the distribution of AUPRC of different methods on two independent test sets, and we find that DISHyper outperforms other advanced methods on both independent test sets. The cancer genes annotated in both OncoKB and ONGene were aggregated from cancer research literature and clinical trials (Chakravarty et al., 2017; Liu et al., 2017), and these cancer genes or data types are not used to train DISHyper. The result suggests that DISHyper has a stronger generalization ability in predicting new cancer genes.

In addition, we illustrate the ability of DISHyper to identify more cancer genes with the analysis of the predicted results. By analyzing the prediction results from different methods, we observed that *WNT5A* is ranked as the top gene (#8) in the DISHyper predictions. In contrast, its ranking is considerably lower in network-based methods (EMOGI and MTGCN), ranks beyond #1500, and even lower in manual feature-based methods (DOGRE and 20/20+), where it ranks beyond #6000. By reviewing the relevant literature, we find that *WNT5A* has been identified as a suppressor gene for breast cancer and is a very potent therapeutic target for breast cancer (Borcherding et al., 2015). Also, *WNT5A* has been identified in several literature and studies as a driver gene in various cancers such as melanoma, prostate cancer, and glioblastoma (Yuzugullu et al., 2009; Radaszkiewicz et al., 2021). Although *WNT5A* is a crucial cancer driver gene, it is only identified as a cancer gene in DISHyper and could not be identified as a cancer gene in any other methods. In addition, we also find many genes with a similar profile to *WNT5A* such as *DKK1* (#22), *SHH* (#23), *FGF10* (#26), *GATA4* (#29), and *TBX1* (#44). These genes are found to be cancer-driver genes or associated with multiple cancers, but these genes are only top-ranked in DISHyper prediction results but are ranked low in other methods. For example, *GATA4* has been identified as an important cancer suppressor gene in lung cancer and may be a potential target for lung cancer therapy (Gao et al., 2019). The result indicates that DISHyper can effectively utilize the knowledge from the annotation gene set to identify cancer genes more accurately. Meanwhile, DISHyper provides a new perspective for cancer gene prediction to reveal those cancer genes that are not discovered by other methods.

### 3.2. DISHyper effectively integrates knowledge from multiple types of annotated gene sets

We integrate multiple types of annotated gene sets in DISHyper, including signaling pathways, GO (Gene Ontology Consortium, 2004), and HPO (Köhler et al., 2021). These different types of annotated gene sets describe the functional associations of genes in different aspects. For example, Signaling pathways contain genes that are jointly involved in a regulatory process or cellular response (Reyna et al., 2020). Integrating knowledge from multiple types of annotated gene sets helps us better characterize the functional association patterns of genes. To illustrate the effectiveness of DISHyper in integrating knowledge from multiple types of annotated gene sets, we evaluate the performance of models trained using data from individual annotated gene sets and their combinations.

As shown in Figure 2C, we find that the performance of the model integrating the three types of annotated gene sets is the best and the performance of the other combinations is also better than using only a single annotated gene set data. The result indicates that there is complementarity between the information of different types of annotated gene sets. Integrating multiple types of annotated gene sets can help us identify cancer genes more accurately. Moreover, the results also illustrate the effectiveness of integrating data from different annotated gene sets based on hypergraphs, and DISHyper may be extended to fuse more data or knowledge.

### 3.3. Characterization of DISHyper-predicted cancer genes by independent functional genomics data

To further illustrate DISHyper’s ability to correctly identify cancer genes, we use cancer genes that have been reported in the literature to analyze the ranking results of DISHyper. We also use independent functional genomics data such as cancer transposons, gene fusions, epigenetic factors, and PPI networks to analyze the characteristics of the DISHyper-predicted cancer gene.

We first use the cancer genes collected in the cancerMine (Lever et al., 2019) database (Download in September 2022) to evaluate our ranking of cancer genes. The cancerMine database is a literature-mining-based cancer gene database that automatically extracts studies of cancer genes in the literature through text-mining tools (Lever et al., 2019). We assess the effectiveness of DISHyper cancer gene rankings by the distribution of annotated cancer genes in each decile of ranking results (Krishnan et al., 2016). We find that cancer genes annotated in cancerMine database are more likely to be ranked high in our prediction results and significantly enriched in the first decile (False Discovery Rate (FDR) = 1.27 × 10^−53^) of the prediction results (Figure 3A). The distribution of cancerMine annotated cancer genes in the DISHyper ranking results shows that the top-ranked genes in the predictions are more likely to be cancer genes. So, we take the top-ranked 200 genes as an example and conduct a series of further analyses of these genes.

**Fig. 3:**
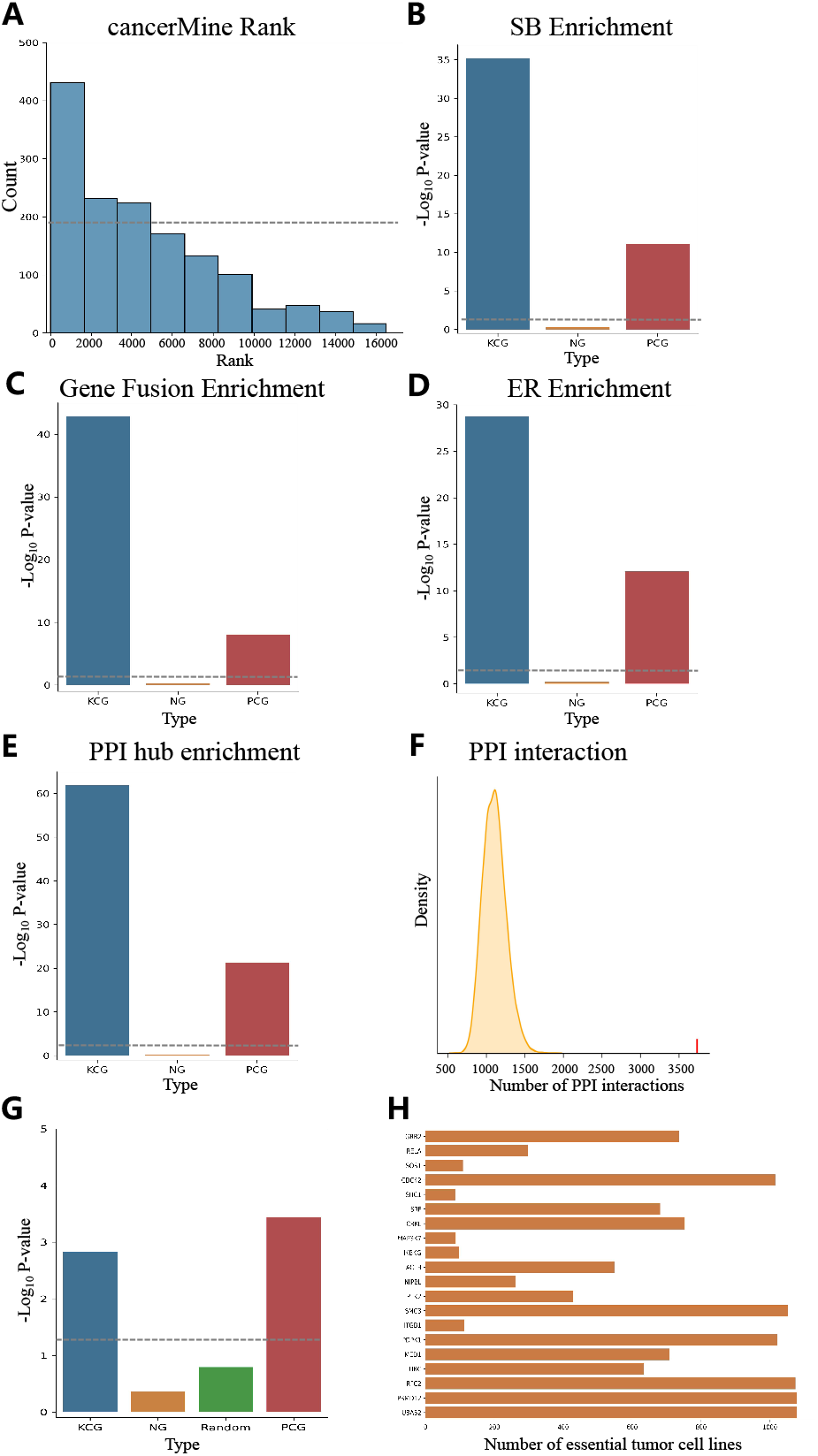
Evaluation of DISHyper-predicted top-ranked 200 cancer genes (PCG) by independent functional genomics data and CRISPR loss-of-function screening results. **(A)** Validation of the ranking results of DISHyper based on the cancerMine database. Fractions of genes (y-axis) indicate the distribution of risk genes in each decile of our predictions. Enrichment analysis of known cancer genes (KCG), neutral genes (NG), and PCG in SB inactivating pattern gene list **(B)**, gene fusion list **(C)**, ER gene list **(D)**, and BioGRID PPI network hub gene list **(E). (F)** Significance analysis of the number of interactions between PCG and KCG on the BioGRID PPI network. The red vertical line indicates the number of interactions between PCG and KCG. The yellow curve indicates the distribution of the interactions between randomly selected genes and KCG. **(G)** Enrichment analysis of KCG, NG, randomly selected genes, and PCG in the essential cancer dependency genes. **(H)** The top-20 PCG with significant negative growth effects on tumor cell lines are displayed in a bar plot. The gray dashline in the figure represents the significant threshold.

Second, we evaluate PCG using Sleeping Beauty (SB) transposon data. The SB insertional mutagenesis is a powerful genetic tool for studying tumor suppressor genes in mammalian cancer models (Dupuy et al., 2009), i.e., screening cancer genes by disrupting gene expression near their insertion sites. We assess PCG using inactivation pattern genes from the Sleeping Beauty Cancer Driver Gene Database (SBCDDB), which collects original mouse models of 19 human tumor types (Newberg et al., 2018). We find that the inactivation pattern genes in the SB transposon study are significantly enriched in known cancer genes (KCG) but not enriched in neutral genes (NG), which is consistent with our expectations (Figure 3B). We also find that inactivation pattern genes are significantly enriched in PCG (*P*-value = 7.43×10^−12^ by Fisher’s exact test ). This suggests that DISHyper can accurately predict potential tumor suppressor genes.

Third, we further evaluate PCG according to cancer fusion gene data. The gene-fusion transcripts and protein products have been considered ideal therapeutic targets and biomarkers for a variety of cancers (Wu et al., 2019). Therefore, we assess the association between DISHyper-predicted PCGs and cancer-related gene fusion events. Gene-fusion events are collected from the TumorFusions (Hu et al., 2018) database and the results of Gao et al (Gao et al., 2018). These two studies examined 33 types of tumor samples from the TCGA project and normal samples in which 20,731 and 25,664 gene fusions were detected, respectively. We find that genes in KCG are significantly enriched in genes that produce oncogenic fusion genes but not enriched in NG (Figure 3C), which suggests that cancer genes are closely associated with gene fusion. Moreover, genes in PCG are also significantly enriched in genes that produce oncogenic fusion genes (*P*-value = 1.31 × 10^−4^, Figure 3C). The result suggests that DISHyper has the ability to predict cancer genes that may work through the gene-fusion mechanism.

Fourth, we explore the possible association between cancer genes and epigenetic regulators (ER). Epigenetic regulators control gene expression through DNA methylation, histone modifications, and chromatin remodeling (Surani et al., 2007). Epigenetic modifications also play a crucial role in cancer. The cancer drugs targeting ER have been extensively studied and applied in the treatment of hematological malignancies. (Cheng et al., 2019). Therefore, we collect a list of 761 ER genes from the EpiFactors database (Medvedeva et al., 2015) and analyze the epigenetic properties of KCG and PCG by this list. We find that ERs are significantly enriched in KCG (*P*-value = 1.83 × 10^−29^) and PCG (*P*-value = 8.13×10^−13^), but not enriched in NG (Figure 3D). The results suggest that epigenetic dysregulation may be a major factor in the influence of these genes on tumor development. Meanwhile, it also shows that DISHyper can discover those cancer genes that affect tumor development through epigenetic modifications.

Finally, we use PPI networks to analyze the network characteristics of DISHyper-predicted PCG. Since the hub genes in the PPI network are more likely to have somatic mutations (Porta-Pardo et al., 2015), we explore the enrichment of cancer genes on BioGRID PPI hub genes. Using 978 genes in the top 5% of the BioGRID PPI network as hub genes, we find that hub genes are enriched in KCG (*P*-value = 1.36×10^−62^) and PCG (*P*-value = 4.75 × 10^−22^), but not enriched in NG (Figure 3E). The result suggests that PCG and KCG have certain traits in common in the PPI network, and these genes play key roles in the PPI network. Many studies have shown that cancer genes are clustered in PPI networks and there are more interactions between cancer genes (Barabási et al., 2011). To assess whether the prediction results of DISHyper would have similar properties, we calculate the significance of the number of interactions between the PCG and KCG in the BioGRID PPI network. Compared with other genes, the PCGs have a significant number of interactions with KCGs (*P*-value *<* 0.0001, Figure 3F). Combining these two results, we find that although DISHyper does not use the information in the PPI network, its prediction results reveal network properties similar to known cancer genes.

### 3.4. DISHyper-predicted cancer genes are essential in tumor cell lines

We use functional validation experiment results to further evaluate DISHyper prediction results. The Broad Institute developed the Cancer Dependency Map (DepMap) database (Tsherniak et al., 2017) and collected the two largest human whole-genome CRISPR screening datasets. Therefore, we use the results from the DepMap database (released on December 22, 2022) to filter a group of essential genes, which significantly affect the survival of multiple cancer cells in CRISPR loss-of-function experiments. We find that PCG has significant enrichment in essential genes (*P*-value = 3.59 × 10^−4^, Fisher exact test, Figure 3G), and KCG also shows significant enrichment, whereas NG and randomly selected genes lack such enrichment. Among the top 20 essential PCG, several genes are found to affect over 1,000 cancer cell lines (Figure 3H). The phenomenon directly raises the question of whether PCG is primarily housekeeping genes that are lethal to any cell when altered. However, this is not the case. We find that only 14% of the genes in PCG affect over 500 cancer cell lines, and most genes only affect fewer than 100 cancer cell lines. Additionally, the KEGG pathway enrichment analysis of PCG shows that PCG is primarily enriched in various cancer pathways (including breast cancer, colorectal cancer, etc., which are not utilized in the training and prediction process of DISHyper), cell differentiation, and Hippo signaling pathway. Taken together, these results indicate that DISHyper not only effectively identifies genes that significantly affect the survival of multiple cancer cells but also that the majority of PCG exhibit cancer-specific cell lethality.

### 3.5. Comprehensive enrichment analysis of DISHyper-predicted cancer genes

We perform enrichment analysis on the DISHyper-predicted cancer genes to gain insights into their biological significance and identify enriched terms. We conduct enrichment analyses on PCG using the GO database, Reactome pathway, and KEGG pathway through the DAVID. In terms of biological processes, PCG are significantly enriched in various positive and negative regulatory activities, such as cell proliferation, cell apoptotic, and gene expression (particularly transcription from RNA polymerase II promoter). Cell proliferation and cell apoptosis are closely associated with the occurrence and development of cancer. In molecular function, PCG are significantly enriched in multiple binding terms, indicating that PCG regulates biological processes by binding with various biomolecules, such as chromatin and proteins. In the Reactome pathway, PCG are primarily enriched in interleukin signaling and diseases related to signal transduction by growth factor receptors. Interleukins and growth factor receptors play crucial roles in tumorigenesis and progression. Additionally, PCG are enriched in multiple kinase signaling pathways, indicating the downstream of these genes may activation of MAP kinases and NF-*κ*B kinases. In the KEGG pathway, PCG significantly enrich in various cancer-related pathways, as well as signaling pathways closely associated with cancer, such as MAPK and Hippo pathways.

### 3.6. Evaluation of the association between novel cancer genes and cancer

We find that 156 genes in PCG have been annotated as cancer genes by the cancerMine database, and the remaining 44 genes are neither in cancerMine nor in NCG and COSMIC CGC. We consider these 44 genes as the novel cancer genes (novelCG). To establish the potential role of novelCG as cancer driver genes, we conducted a comprehensive analysis using cancer sample data from the TCGA study.

We first analyze novelCG based on the OncoDB database, which integrates RNA-seq, DNA methylation, and clinical data from more than 10,000 tumor patients in the TCGA study and normal samples in the GTEx study (Tang et al., 2022). Among the 44 novelCG, 41 genes exhibit significant differential expression in one or more cancer types (FDR-adjusted *P*-value (*Q* -value) *<* 0.05 and |*log*2*FC*| *>* 1), and 31 genes are significantly differentially methylated in one or more cancer types (*Q <* 0.05 and |*Beta*| *>* 0.2). Furthermore, in the analysis of clinical characteristics andpathological diagnostic phase, we find that the expressions of 37 genes are significantly associated with the size and extent of the primary tumor (Pathological T stage, *Q <* 0.05), and 28 genes are significantly associated with the distal spread of one or more tumors (Pathological M stage, *Q <* 0.05).

Then, we perform survival analysis of novelCG on multiple cancer types based on the GEPIA2 platform, which collected cancer samples and normal samples from the TCGA study and GTEx study and generated the results of survival analysis of genes in different cancer types by RNA sequencing data (Tang et al., 2019). With this survival map, we find that 43 of 44 novelCG genes have significant survival analysis results in multiple cancer types (*P*-value *<* 0.05), and this result indicates that novelCG expression significantly affects the prognostic outcome of multiple cancer types. We also find that *TREX1* dose not produce significant associations with cancer in terms of differential expression, differential methylation, or pathological diagnostic stage, but the gene is significantly associated with CESC, HNSC, and two other cancers in the survival analysis. These findings suggest that these 44 novelCG genes may influence tumorigenesis and progression in diverse ways.

Additionally, there is supporting evidence from many aspects to support our predictions. We find that 35 genes in novelCG may generate oncogenic fusion genes, indicating that these genes potentially influence the occurrence and progression of various tumors through gene fusion mechanisms. In the CRISPR loss-of-function experiments results, We find that nine genes in novelCG significantly impact multiple cancer cell lines, and these genes exhibit cancer cell-specific lethality. We also observe that 20 genes in novelCG significantly impact the proliferation of cell lines associated with one or more common cancer types. Meanwhile, novelCG exhibits numerous interactions with known cancer genes in the STRING PPI network. Furthermore, we conduct KEGG pathway enrichment analysis for novelCG, revealing significant enrichment in Hippo and TGF-*β* signaling pathways. These pathways are closely associated with cancer, highlighting the research potential of novelCG. The above results demonstrate the reliability of our predictions and suggest that novelCG likely includes potential cancer genes.

## 4. Discussion

In this paper, we introduce DISHyper, a novel hypergraph-based cancer gene prediction method. DISHyper extracts higher-order gene association information from the annotated gene set by the hypergraph neural network to identify new cancer genes. DISHyper is different from the previous methods in principle. Methods such as 20/20+, DORGE, and EMOGI identify cancer genes based on mutational, epigenetic, or biological network data. Although these methods have achieved certain success, they all focus on a single feature of cancer genes and have difficulty in describing the complex role of cancer genes in tumor development. DISHyper not only significantly outperforms other methods on both pan-cancer datasets and several independent test sets, but also can reveal cancer genes that cannot be identified by other methods, such as shown in our study. The above results strongly indicate that DISHyper can identify cancer genes more accurately and reliably by integrating expert and domain knowledge from annotated gene sets.

In summary, this work highlights the integration of annotated gene set data based on hypergraphs to achieve more comprehensive and accurate predictions of cancer genes. DISHyper will be an essential resource for cancer genetic research and a very significant breakthrough for the study of cancer gene prediction methods. DISHyper still has space for further improvement. The DISHyper only uses annotated gene set data, and the fusion of mutation, epigenetic, and biological network data on the basis of DISHyper may further improve the performance of the model. In future work, we plan to fuse more data in DISHyper and conduct further studies on specific cancer types such as breast cancer and lung cancer.

## 6. Acknowledgments

This work was supported in part by the National Key Research and Development Program of China (No. 2021YFF1201200). This work was carried out in part using computing resources at the High-Performance Computing Center of Central South University.

